# The P600 during sentence reading predicts behavioral and neural markers of recognition memory

**DOI:** 10.1101/2024.03.08.584041

**Authors:** Friederike Contier, Melissa Höger, Milena Rabovsky

## Abstract

The P600 ERP component is elicited by a wide range of anomalies and ambiguities during sentence comprehension and remains important for neurocognitive models of language processing. It has been proposed that the P600 is a more domain-general component, signaling phasic norepinephrine release from the locus coeruleus in response to salient stimuli that require attention and behavioral adaptation. Since such norepinephrine release promotes explicit memory formation, we here investigated whether the P600 during sentence reading (encoding) is thus predictive of such explicit memory formation using a subsequent old/new word recognition task. Indeed, the P600 amplitude during our encoding task was related to behavioral recognition effects in the memory task on a trial-by-trial basis, though only for one type of violation. Recognition performance was better for semantically, but not syntactically violated words that had previously elicited a larger P600. However, the P600 to both types of violations during encoding was positively related to a more subtle, neural marker of recognition, namely the amplitude of the recollection ERP component in response to old words. In sum, we find that the P600 predicts later recognition memory both on the behavioral and neural level. Such explicit memory effects further link the late positivity to norepinephrine activity, suggesting a more domain-general nature of the component. The connection between the P600 and later recognition indicates that the neurocognitive processes that deal with salient and anomalous aspects in the linguistic input in the moment will also be involved in keeping this event available for later recognition.

## Introduction

Language comprehension is a multifaceted cognitive process that involves the rapid evaluation and integration of different kinds of information. In the neural signature, one facet of this process is characterized by the P600, a component of the event-related potential (ERP) typically observed between 500 and 900 ms after the onset of a critical word or phrase. The P600 is best known to be sensitive to syntactic ambiguities and violations (e.g., Friederici et al., 1993, 1996; Gouvea et al., 2010; Münte et al., 1998; Osterhout & Mobley, 1995) and traditionally implicated as a marker of structural processing, such as re-analyses, integration, or effort required to build a coherent sentence structure (Friederici, 1995, 2002; Gouvea et al., 2010; Hagoort, 2003; Kaan et al., 2000; Osterhout et al., 1994). However, accumulating evidence suggests that the P600 is sensitive to a wide range of linguistic anomalies, for instance, regarding semantics, pragmatics, and even spelling (e.g., Brothers et al., 2020; Hoeks et al., 2004; Kim & Osterhout, 2005; Kuperberg, Holcomb, et al., 2003; Münte et al., 1998; Regel et al., 2014; van de Meerendonk et al., 2011). More recent theories thus view the P600 as a more general signal of error monitoring or correction in language (Ryskin et al., 2021; van de Meerendonk et al., 2009, 2010) or difficulty to establish a single, coherent utterance representation (Brouwer et al., 2012; Brouwer & Hoeks, 2013; Li & Ettinger, 2023). The neurocognitive foundation of the P600, as well as its connections with other cognitive processes and domains, thus continue to be areas of ongoing uncertainty.

Instead of reflecting a process specific to language as discussed above, the P600 could alternatively be part of a family of domain general, late positivities (Coulson et al., 1998a, 1998b; Gunter et al., 1997). Anomalies eliciting a P600 are not restricted to language as they also involve, for instance, incongruencies in music, within non-linguistic sequences, arithmetic rules, and narrative structures in comics, scenes and events (e.g., Christiansen et al., 2012; Cohn et al., 2014; Cohn & Kutas, 2015; McLean et al., 2020; Núñez-Peña & Honrubia-Serrano, 2004; Patel et al., 1998; Sitnikova et al., 2008; Võ & Wolfe, 2013). A response to such a variety of salient, incongruent stimuli is reminiscent of the P3(b). The P3 is a widely studied ERP component, which is most sensitive to ‘oddballs’, that is, rare and relevant target stimuli among frequent standard stimuli across domains (Donchin, 1981). Indeed, the P3 and P600 are both late, positive-going components and the morphology of their waveforms seem indistinguishable (Sassenhagen & Fiebach, 2019). Both components are most sensitive to stimulus probability and task relevance (see Leckey & Federmeier, 2019, for a review) and require attention and cognitive control (Brothers et al., 2021; Contier et al., 2022; Gratton et al., 2018; Xu et al., 2021). P3 latency is highly dependent on stimulus and task complexity (Kutas et al., 1977) which would explain latencies of 600-700 ms on complex linguistic stimuli, such as anomalies in sentences.

Besides the temporal difference between the P600 and P3, there is further evidence potentially questioning the hypothesis that the two components are related. For instance, physical anomalies and linguistic agreement errors together lead to a larger positivity than either anomaly by itself (Osterhout et al., 1996). This could indicate that two independent components (P3 and P600) “add up” (Osterhout et al., 1996), but alternatively, that a double anomaly simply makes the stimuli more salient, which is known to increase P3 amplitude (Coulson et al., 1998a; see also Leckey & Federmeier, 2019). Furthermore, an essential premise of the “P600-as-P3” hypothesis is that the two components have a common neural generator. However, several lesion studies (Frisch et al., 2003; Hagoort et al., 2003) indicate that essential neural substrate for the generation of the P600 might be fundamentally different from what has been found for the P3 (see e.g., Linden, 2005; Soltani & Knight, 2000). This mixed evidence has led some researchers to reject the hypothesis (e.g., Friederici et al., 2001; Frisch et al., 2003; Osterhout et al., 1996; Osterhout & Hagoort, 1999). Therefore, despite many similarities, the debate about the “P600-as-P3” hypothesis has not yet settled and requires further investigation. If the P600 is indeed related to the P3, this would indicate that some processing aspects of sentence comprehension are not unique to language, but are shared with other domains.

It has been specifically proposed that both the P3 and P600 reflect phasic norepinephrine (NE) release from the locus coeruleus (LC) as a general response to salient and motivationally significant events (Bornkessel-Schlesewsky & Schlesewsky, 2019; Murphy et al., 2011; Nieuwenhuis et al., 2005; Sassenhagen et al., 2014; Sassenhagen & Bornkessel-Schlesewsky, 2015; Vazey et al., 2018). At target cortices, NE leads to more focused attention to relevant stimuli (neural gain, Nieuwenhuis et al., 2005) or acts as an interrupt signal (neural network reset, Sara & Bouret, 2012). Mirroring the effect of infrequent stimuli on the P3 and P600, there is evidence for a strong oddball effect on the phasic LC/NE activity (e.g., Aston-Jones et al., 1997), which is also highly dependent on task relevance and attention (Nieuwenhuis et al., 2005). One important piece of evidence linking the positivities to NE is that in contrast to other ERPs which are strictly onset-aligned, the latency of both the P3 (e.g., Kutas et al., 1977; Verleger et al., 2005) and P600 (Sassenhagen et al., 2014; Sassenhagen & Bornkessel-Schlesewsky, 2015) is more aligned to the behavioral response, which is a crucial feature of the LC/NE response (Bouret & Sara, 2004). Further, the amplitude of both components covaries with markers of NE, such as the task-evoked pupillary response (Contier et al., 2024). Beyond correlational evidence, intervention studies (e.g., Joseph & Sitaram, 1989; Pineda & Westerfield, 1993; Rufener et al., 2018; Ventura-Bort et al., 2018) as well as lesion studies in non-human animals (e.g., Ehlers & Chaplin, 1992; Pineda et al., 1989) in fact suggest a causal relationship between the P3 and norepinephrine. Based on our knowledge of LC/NE activity, specific predictions can be made about the behavior of the P3 and P600 and their downstream effects, for instance with regards to their link to memory.

The LC/NE system plays a crucial role in explicit memory formation (Gibbs et al., 2010; Sara, 2015). Norepinephrine is proposed to enhance the responsiveness of cortical neurons to incoming signals and suppress background noise, making it easier to detect and process relevant stimuli (Aston-Jones & Cohen, 2005). This improved processing can in turn facilitate encoding and consolidation of explicit memories. Furthermore, the LC concurrently sends noradrenergic projections to limbic structures crucial for memory encoding and retrieval, such as the hippocampus, amygdala, and entorhinal cortex (e.g., Hansen & Manahan-Vaughan, 2015; Harley, 2007; McGaugh, 2000; Roozendaal et al., 2009; Schutsky et al., 2011). Indeed, both correlational studies and studies manipulating NE in humans and non-human animals suggest that more NE release enhances explicit memory (e.g., Clewett et al., 2018; Devauges & Sara, 1991; Gold & Van Buskirk, 1975; Hämmerer et al., 2018; Hauser et al., 2019; Schutsky et al., 2011). Thus, if the late positivities reflect LC/NE activity and LC/NE activity also enhances explicit memory formation, the positivities themselves should also relate to such explicit memory formation.

Indeed, late positive ERPs during encoding have been linked to memory formation. Generally, rare and deviant stimuli that stand out (‘isolates’) are better remembered than the rest of the stimuli (von Restorff, 1933). This ‘von Restorff effect’ thus links the oddball effect to memory. Crucially then, within such isolates, the amplitude of the P3 amplitude during processing (encoding) is predictive of later memory of those stimuli: Stimuli that elicit a larger P3 at encoding are more likely recognized and/or recalled later (Azizian & Polich, 2007; Dunn et al., 1998; Fabiani et al., 1986, 1990; Fabiani & Donchin, 1995; Kamp et al., 2013, 2015, 2012; Kamp & Donchin, 2015; Karis et al., 1984; Paller et al., 1988; Sanquist et al., 1980; Voss & Paller, 2009). These so-called subsequent memory effects (Mecklinger & Kamp, 2023) are highly variable in their latency and are thus often just referred to as late positive potentials (e.g., Paller et al., 1988). These effects are robust across lower and higher level stimulus categories (e.g., Fabiani & Donchin, 1995; Kamp et al., 2013; Otten & Donchin, 2000) and emotionally salient isolates (Kamp et al., 2015) as well as for both recognition and free recall (e.g., Fabiani & Donchin, 1995; Kamp & Donchin, 2015; Karis et al., 1984; Sanquist et al., 1980). Such a predictive relationship between a late positivity during encoding and subsequent memory has also been found in more complex, linguistic and semantic contexts (e.g., Friedman & Trott, 2000; Höltje et al., 2019; Neville et al., 1986; Paller et al., 1987, 1988). For example, Neville and colleagues (1986) showed participants a categorization prompt (‘*A type of bird.’*) to which the subsequently presented completion was either semantically congruent (‘*Robin*’) or incongruent (‘*Nail*’). Completions that elicited a larger late positivity (between 400-950 ms) were better recognized in a subsequent old/new memory test. Despite this strong evidence associating late, positive ERP components like the P3 with explicit memory formation, and the observations of similar effects within complex semantic tasks where the latency corresponds to the P600, it remains uncertain whether the typically observed P600 during sentence comprehension is also directly tied to explicit memory processes, given the lack of studies within standard P600 task paradigms.

Here we investigated how the P600 relates to explicit memory. We used a classic sentence reading paradigm to elicit the P600 in conjunction with a subsequent recognition task to measure explicit memory. If the P600 reflects norepinephrine release and norepinephrine is involved in explicit memory formation, the amplitude of the P600 should positively predict whether the respective target word is later correctly recognized in an old/new word decision task. We investigated whether these memory effects arise as a consequence of saliency due to semantic and (morpho-)syntactic anomalies. Besides behavioral recognition^1^, we also explored whether the P600 is related to a neural, more fine-grained measure of memory. Specifically, we explored whether the P600 is related to the amplitude of the parietal old/new recollection ERP component that has been proposed to respond to the amount of information explicitly recollected (Curran et al., 2006; Friedman & Johnson, 2000; Rugg & Curran, 2007).

## Method

Sample size, hypotheses, and respective statistical models were pre-registered on the Open Science Framework (https://osf.io/pv964/?view_only=e8b52f2a19144ca7aee878491c0c3836). Any deviations from this plan are explicitly stated. The associated OSF project also hosts all pre-processing and analyses scripts as well as the pre-processed data.

### Participants

As preregistered, the sample included 36^2^ German native speakers (28 female; mean age: 23.4 years, range: 18-38) with normal or corrected vision. None of them reported any history of neurological or psychiatric disorder or language impairments. Participants were right handed, as assessed using the Edinburgh Handedness Inventory (Oldfield, 1971; 12-item version; 82.8, range: 50-100). Participation was compensated monetarily or with course credit. Six participants had to be replaced, because they were either uncooperative (1), ambidextrous (1), or misunderstood the memory task (4). Each individual provided written informed consent. The experiment was part of a research project whose protocols were approved by the Ethics Committee of the German Psychological Association (DGPs; MR102018).

### Procedure & materials

Each experimental session started with the encoding task (sentence comprehension), followed by a 5-minute break and the recognition task. Participants were not informed about the memory test (recognition task) until it started since we were interested in incidental memory effects and tried to avoid strategic memorizing that would influence “natural” processing of the sentences (Meyer et al., 2007). Participants were seated in an acoustically and electrically shielded booth. Presentation in both tasks was controlled via the *Psychophysics Toolbox* (Brainard, 1997) in *MATLAB* R2020a (The_Mathwork_Inc., 2020).

Stimuli were presented visually on a computer screen with black, Geneva font on grey background (see Fig. 1) at 1.415° visual angle.

**Figure 1.**
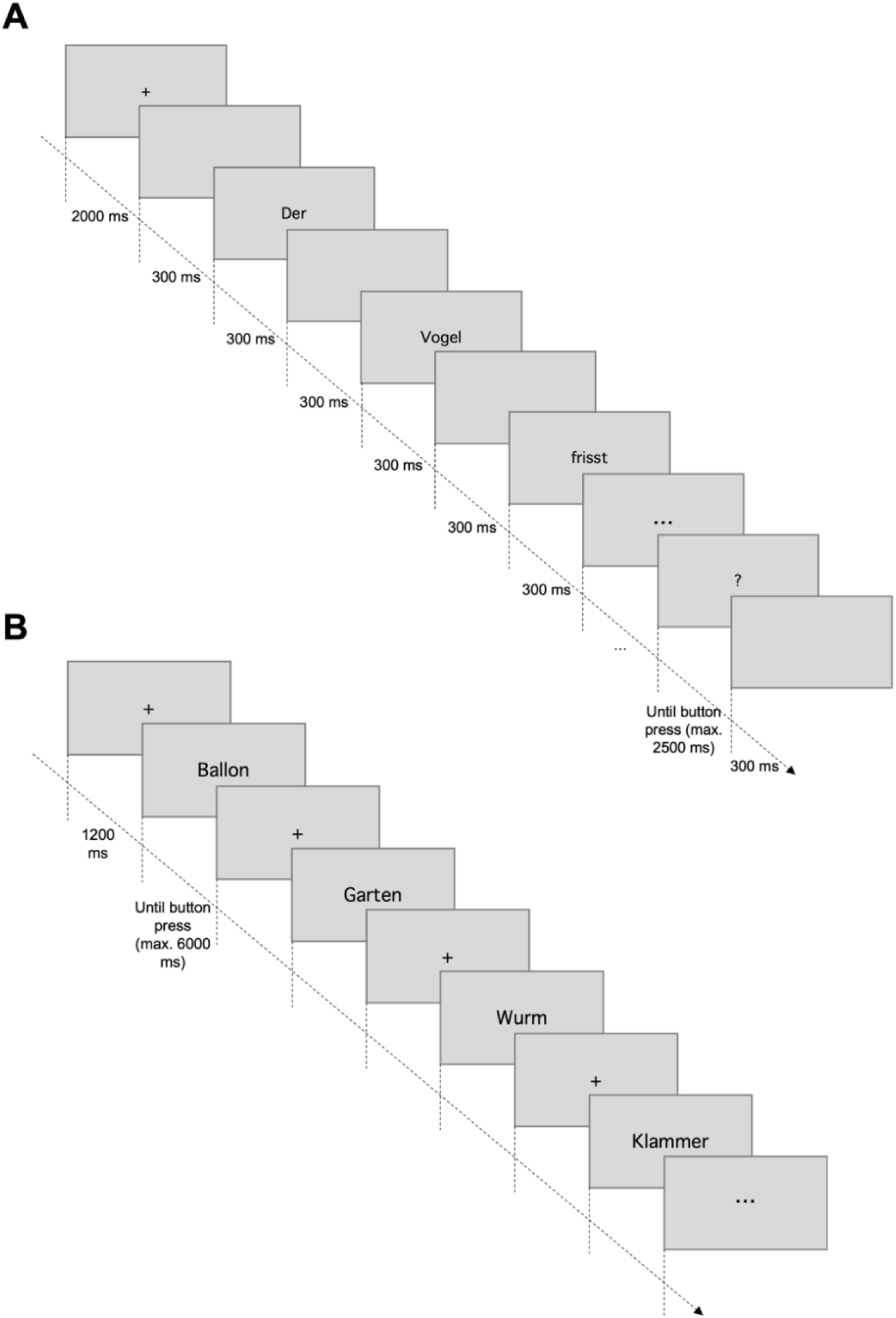
**A**: Word-by-word sentence presentation of one trial in the encoding task. The question mark appeared after each sentence and prompted the participant to respond via button press whether the preceding sentence was anomalous or not. **B**: Stimulus presentation in the recognition task. Participants were instructed to respond whether the respective word had appeared as part of the sentences in the first task (i.e., encoding task) or not, starting as soon as the word appeared.

#### Encoding task

The encoding task was a classic sentence comprehension task designed to elicit the P600. Participants were visually presented with sentences that were either correct control sentences or contained a morphosyntactic (for simplicity, henceforth called “syntactic”) or semantic violation. Violation sentences were constructed from a set of 208 well-formed and meaningful German sentences (e.g., *Die Wolke verdeckt den <u>Mond</u> komplett*. “The cloud covers the <u>moon</u> completely.”). Syntactic violations consisted of agreement violations between the gender of the target noun and its determiner (\**Die Wolke verdeckt die_sg.fem._ <u>Mond_sg.masc_</u>. komplett.* “The cloud covers the_sg.fem._ <u>moon_sg.masc._</u> completely”). Semantic violations were created by exchanging the target noun with a target noun from another sentence in the set that would create the most implausible sentence meaning (\**Die Wolke verdeckt das <u>Interview</u> komplett.* “The cloud covers the <u>interview</u> completely”). Sentence lengths varied between five and ten words. Target words were thus always nouns, and their position within the sentence varied between mid-sentence and final word position.

Each of the 208 experimental sentences thus had three versions: one correct control, one syntactic violation, and one semantic violation version. Importantly, this also entails that, across participants, each target word appeared in all three conditions. Following latin-square design, we created four lists, each including 104 of the sentences in their control version, another 52 in their syntactic violation and the remaining 52 in their semantic violation version. Thus, each experimental sentence appeared in one list with a syntactic violation, in one list with a semantic violation, and in two lists as a correct control sentence. In order to decrease the proportion of violations without further complicating this counterbalancing procedure, we added (the same) 32 correct filler sentences to each list. Including the fillers, which were not part of the analysis, each participant was thus presented with 240 sentences with a violation proportion of 22% for each violation type.

Our aim was further to decrease variability in temporal lag between encoding and retrieval between stimuli. Thus, sentences in each set were further divided into two blocks. The same was done with the word stimuli of the subsequent recognition task (see below), matching the assignment of the respective words to that from the sentence comprehension task. Sentences were randomly pre-assigned to a block, but each block contained the same proportion of each stimulus type described above. Order of blocks was counterbalanced across participants and kept constant between the sentence reading and the recognition task, thus resulting in eight stimuli lists (4 sentence lists x 2 block orders). For each participant, sentences within each block were presented in random order with the constraint of no consecutive violation sentences of the same type. For the subsequent recognition task, the words were randomized within each block independently from the order in the sentence comprehension task. Note that despite this parallel blocking procedure, the recognition task did not start until the entire encoding task was finished.

As depicted in Figure 1A, each trial started with a fixation cross presented for 2000 ms. Then, the sentence was presented word by word in the center of the screen for 300 ms per word with a 300 ms blank inter-word interval. The presentation of non-target words was prolonged for words with more than eleven letters by 16.167 ms (one screen presentation frame rate) per additional letter (for comparable approaches, see, e.g., Hodapp & Rabovsky, 2021; Nicenboim et al., 2020). 300 ms after the offset of the last word, a question mark appeared, prompting the participant to judge as accurately and quickly as possible whether the preceding sentence was correct or (grammatically or semantically) anomalous. Two marked control keys on the keyboard served as response buttons, pressed by the left and right index finger, respectively. Stimulus-response mapping was counterbalanced across participants. The question mark was shown until button press, but for a maximum of 2500 ms. The inter-trial interval lasted 300 ms.

Participant-controlled rest breaks were provided every 40 trials. Before experimental sentences were presented, participants were given written instructions and practiced the task with four example sentences. The task lasted approximately 40 minutes.

#### Recognition task

The old/new word recognition task served as a measure of how well target words from the preceding sentences were explicitly remembered. The condition “seen” comprised all 208 presented target nouns from the experimental sentences in the preceding encoding task. As “unseen” distractors, we selected 208 additional nouns from the SUBTLEX-DE database (Brybaert et al., 2011). Since our priority were the responses to the “seen” words, not the old/new effect per se, our priority was that “unseen” distractors should not distort the memory of the “seen” words (i.e., adding more variance in their recognition, unrelated to encoding). Thus, we specifically selected nouns that would not have fit semantically into any of the sentences in the encoding task (i.e., in place of the target word), thereby possibly creating false retrospective memory and adding uncertainty about the actual target word of that sentence. This was specifically necessary as 22% of seen words appeared as a semantic mismatch in their sentences, thus a semantically fitting alternative might match the memory trace for a (expected but not seen) congruous continuation (Hubbard et al., 2019; Rommers & Federmeier, 2018). Two independent raters judged the semantic mismatch of these nouns into all experimental sentences in the encoding task using a 3-point scale (1= doesn’t fit into any sentence, 3 = fits into several well). The two raters agreed on 80% of the cases and in cases of disagreement, a third rater decided the case. Final words had an overall mean rating of 1.226 (with no ratings of 3). We still aimed to keep linguistic characteristics^3^ between “seen” and “unseen” words as similar as possible: The two word types did not significantly differ in length (seen *M* = 7.029 letters, *SD* = 2.711; unseen *M* = 7.024 letters, *SD* = 2.272; *t*(420) = - 0.029, *p* = .977) or frequency (seen *M* = 20.22, *SD* = 68.225; Unseen *M* = 15.41, *SD* = 26.816; *t*(411) = 0.987, *p* = .324). Only neighborhood size was significantly larger for seen (*M* = 13.46, *SD* = 17.918) than unseen words (*M* = 9.923, *SD* = 11.865; *t*(411) = 23.944, *p* = .017). However, note that this means there are slight differences in stimulus characteristics between “seen” and “unseen” words which is relevant for experimental effects between the two conditions (such as the old/new ERP effect). However, of primary interest for the present study were behavioral and ERP responses within the “seen” category only and how they relate to ERPs at encoding. As noted above, “seen” words appeared in the same of two blocks in which they also appeared in the encoding task to reduce variance in delay between encoding and recognition between stimuli. However, the words were presented randomly within each block independently from the order in the sentence comprehension task.

As displayed in Figure 1B, each trial started with a fixation cross, presented for 1500 ms. Then, the word was presented at the center of the screen until the participant’s response or 6000 ms had passed. Participants were instructed to judge as accurately as possible whether the respective word had appeared in one of the sentences in the preceding task (encoding task) or not. Two marked (control) keys on the keyboard served as response buttons, pressed by the left and right index finger, respectively. Stimulus-response mapping was counterbalanced across participants. However, the mapping was constant across the two tasks: Conservatively, the “sentence correct” button in the encoding task always served as the “seen” button in the recognition task. The inter-trial interval lasted 1500 ms.

Participant-controlled rest breaks were provided every 45 trials. Before the start of the experimental trials, participants were given written instructions. The task lasted approximately 25 minutes.

### EEG data acquisition and processing

In both tasks, we recorded participants’ electroencephalogram (EEG) using 32 passive Ag/AgCl electrodes (actiCHamp Plus system, Brain Products GmbH, Gilching, Germany), spaced according to the international 10-20 system. Impedances were kept below 5 kΩ. The ground electrode was positioned at FCz and the reference at the right mastoid. Bilateral VEOG as well as right HEOG electrodes were in place to capture eye movements and blinks. The EEG was recorded at sampling rate of 1000 Hz (bandpass filter of .016-250 Hz, time constant of 10 s).

We preprocessed the data in *MATLAB* R2020a (The_Mathwork_Inc., 2020), using the *EEGLAB* toolbox (Delorme & Makeig, 2004). First, we downsampled the data to 500 Hz and re-referenced to the average of the left & right mastoid.

Independent component analysis (Infomax ICA, Jung et al., 2001) was used to correct ocular artifacts. ICA was run on specifically prepared continuous data which was filtered using a 1-30 Hz Butterworth filter, epoched from −1 to +2s post target word onset, and cleaned of extensive artifacts (removal of epochs exceeding +150/-150 mV; see e.g., Kappenman et al., 2021). The *IClabel* plug-in (Pion-Tonachini et al., 2019) then automatically categorized independent components (ICs) based on their assumed source. We removed independent components identified to be eye-related with a probability > 30% (number of excluded ICs in encoding task: *M* = 3.34, *SD* = 0.99 and recognition task: *M* = 3.56, *SD* = 0.81) from the downsampled and re-referenced data. We additionally removed (max. three) channel noise components based on visual inspection.

Corrected, continuous data were high-pass filtered (0.1 Hz, two-pass Butterworth with a 12 dB/oct roll-off) and low-pass filtered (30 Hz, two-pass Butterworth with a 24 dB/oct roll-off). Then, we interpolated bad channels (z-score larger than +/- 3) using a spherical spline function and epoched data from −200 to 1100 ms, time-locked to target word onset. Finally, we baseline corrected epochs relative to the 200 ms interval preceding the onset of the target word. Artifactual epochs exceeding +/-75 mV were automatically removed.

In the encoding task, we additionally excluded EEG epochs with incorrect responses. On average, after data exclusion participants provided 48 syntactic violation sentence trials (range: 38-51), 43 semantic violation sentence trials (range: 23-50), and 93 control sentence trials (range: 80-103) in the encoding task. In the recognition task, participants provided ERP data from 191 “seen” trials (range: 165-202) and 193 “unseen” trials (range: 178-205). Note that, depending on the analysis, number of trials might deviate from this summary as we tried to include as many trials as feasible in each scenario. For example, analyses testing the relationship between encoding P600 and recognition performance includes all trials from the encoding task that had not been rejected (due to EEG artifacts or incorrect responses) and respective behavioral data from “seen” trials in the recognition task (irrespective of EEG artifacts in these trials). However, the analysis relating the encoding P600 to the recognition ERP naturally only includes the trials without artifacts in either of the tasks.

The P600 on semantic and syntactic violations in the encoding task was quantified as the mean amplitude 600-900 ms after target-word onset within a parietal region of interest (ROI: CP1, CPz, CP2, P3, Pz, P4, PO3, POz, PO4; e.g., Contier et al., 2022; Sassenhagen & Bornkessel-Schlesewsky, 2015; Tanner et al., 2017). For exploratory analyses, we also extracted N400 values in the control and semantic condition of the encoding task, quantified as the mean amplitude 300-500 ms after target word onset within a centro-parietal channel cluster (ROI: Cz, CP1, CPz, CP2, Pz; see e.g., Hodapp & Rabovsky, 2021; Kuperberg et al., 2020). In the recognition task, the parietal old/new recollection ERP component was quantified as the mean amplitude 500-800 ms after word onset within a left-parietal channel cluster (ROI: C3, Cz, CP5, CP1, CPz, P3, Pz, PO3, POz; Curran et al., 2006; Friedman & Johnson, 2000; Rugg & Curran, 2007; Wilding & Ranganath, 2011). As preregistered, we additionally explored the mid-frontal old/new ERP component (FN400) as a measure of familiarity and/or implicit memory (Curran et al., 2006; Mecklinger, 2006; Rugg & Curran, 2007; Wilding & Ranganath, 2011). This FN400 was calculated as the mean amplitude 300-500 ms after word onset within a mid-frontal channel cluster (ROI: AFz, F3, Fz, F4, FC1, FC2; see e.g., Curran et al., 2006; Rugg & Curran, 2007; Wilding & Ranganath, 2011).

### Statistical analyses

To account for subject and item level variability, we performed mixed-effects model (LMM) analyses using the package *lme4* (Bates et al., 2014) as implemented in *R* (R core Team, 2018). All models included random slopes by participant and item (sentence / word) for the predictor(s) of interest, except in cases where the predictor was not manipulated within subjects or items. Following the recommendations by Barr et al. (2013), we tried to fit this maximal random effect structure and reduced its complexity successively until the model converged. We did so by increasing the number of iterations to 10 000 and introducing an optimizer (“bobyca”), then removing correlations between random intercepts and slopes and if necessary, entire random slopes. Predictors with two levels of interest were sum coded (- .5/.5, e.g., violations vs. controls), whereas for predictors in which the second level had several sublevels of interest (e.g., semantic vs. syntactic violations), Helmert coding was used (Brehm & Alday, 2022; Schad et al., 2018). The significance of fixed effects was determined via likelihood ratio tests that compared the fit of the model to that of a null model lacking the respective fixed effect but including all remaining fixed effects as well as the identical random effect structure. Group contrasts were calculated using the package *emmeans* (Lenth et al., 2023). We used the standard *p* < .05 criterion for determining if the likelihood ratio tests suggest that the results are significantly different from those expected if the null hypothesis were correct. Response times (RTs) were log-transformed, as they were not normally distributed (mean skewness in encoding task: 1.717, *SE* = 0.03; recognition task: 2.258, *SE* = 0.02). Plots were generated in *MATLAB* (The_Mathwork_Inc., 2020) and *R* (ggdist, Kay & Wiernick, Brenton, 2023; sjPLot, Lüdecke, 2023; ggforce, Pedersen, 2022; gghalves, Tiedemann, 2022; ggplot2, Wickham, 2009).

## Results

### Sentence processing task

#### Behavioral performance

Accuracy was generally good (*M* = 91%, *SD* = 4), indicating that participants focused on the task. Performance did not differ between syntactic violations (*M* = 95%, *SD* = 4) and correct control sentences (*M* = 95%, *SD* = 4; *β* = −0.069, *SE* = 0.124, *z* = −0.559, *p* = .576). However, semantic violations were identified significantly worse (*M* = 83%, *SD* = 13) than correct and sensible control sentences (*β* = −1.059, *SE* = 0.188, *z* = −5.619, *p* < .001).

#### Sentence ERP effects

Mean ERP amplitudes over time in each of the three conditions are shown in Figure 2. Overall, we found a P600 effect: Amplitudes between 600-900 ms at parietal sites were significantly larger on violations than correct controls (*β* = −2.402, *SE* = 0.352, *χ2* = 31.318, *p* < .001). As expected, this P600 effect was significantly larger regarding syntactic violations than semantic violations (*β* = 3.411, *SE* = 0.411, *χ2* = 40.666, *p* < .001). To check whether there was a semantic P600 at all, an additional model directly tested each type of P600 (i.e., compared the amplitude for each violation type directly to the amplitude on controls). This model confirmed the syntactic P600 (*β* = 4.123, *SE* = 0.411, *χ2* = 48.332, *p* < .001). However, the difference between the semantic violations and correct controls at these parietal channels was only a positive trend (*β* = 0.708, *SE* = 0.389, *χ2* = 3.256, *p* = .071). The topographic plot in Figure 2 indicates that the effect was stronger at left-parietal and frontal channels, in congruence with previous reports (Kuperberg et al., 2020; Stone et al., 2023). Indeed, post-hoc analyses confirmed that amplitudes were larger for semantic violations than correct controls at both a left-parietal (channels P3, CP5, P7; *β* = 0.989, *SE* = 0.353, *χ2* = 7.313, *p* = .007) as well as a frontal (channels AFz, Fz, FC1; *β* = 0.946, *SE* = 0.429, *χ2* = 4.705, *p* = .03) cluster. We thus additionally used this combined left-frontal cluster in subsequent analyses as a measure of the semantic P600 in particular.

**Figure 2.**
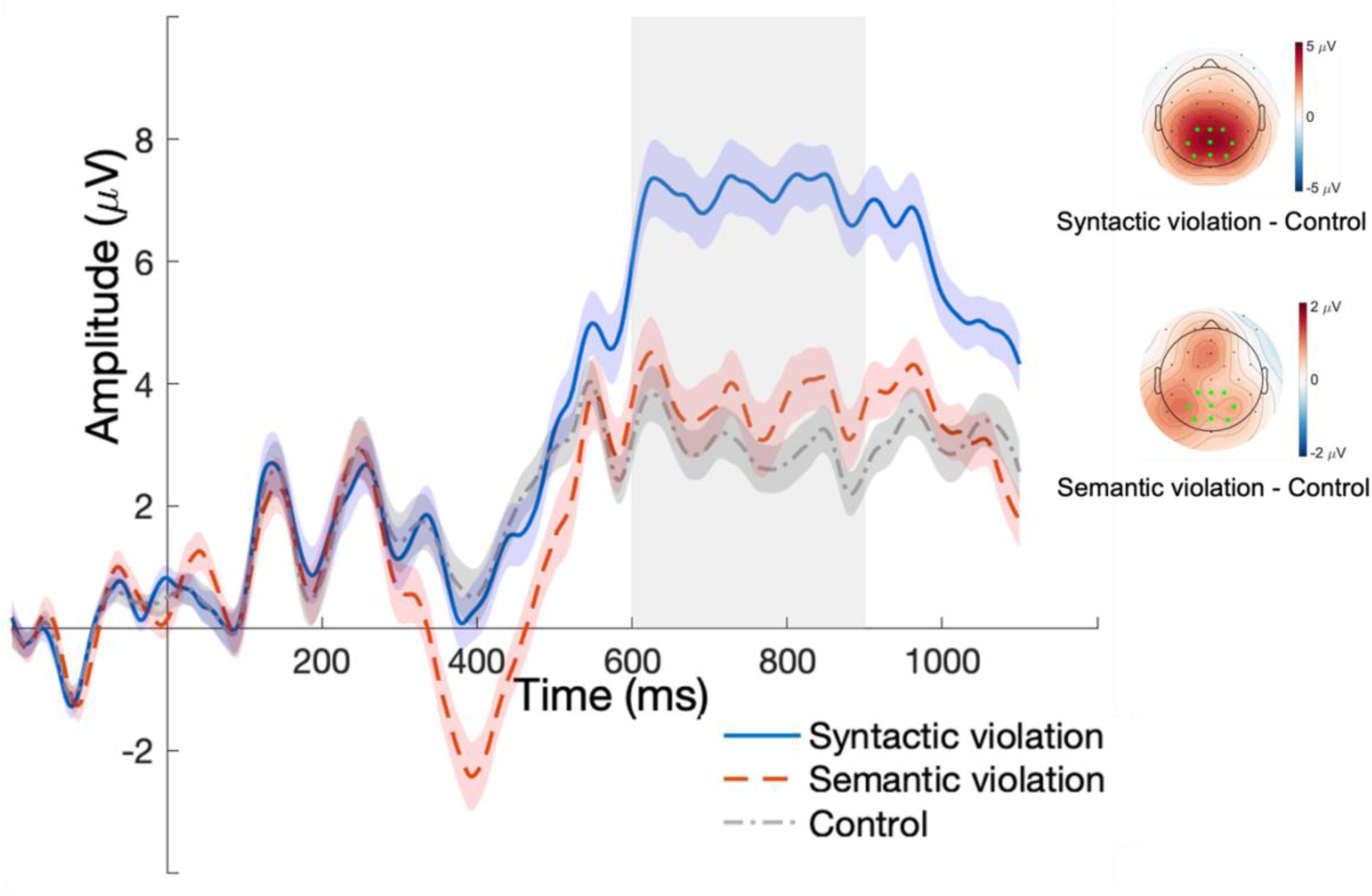
Grand averaged ERP amplitude for the three conditions of the sentence processing task. Time series are locked to the onset of the target word within the posterior ROI channel cluster (marked in light green on the topographic maps). Trial-wise P600 amplitudes were calculated as the mean in the time interval marked by the grey box (600-900 ms). The topographic maps show the mean amplitude difference between violations of each type and controls within the same time window. Error bands indicate the standard error of the mean (SEM).

An additional, exploratory model also identified an N400 effect: Amplitudes between 300-500 ms at the centro-parietal cluster were significantly more negative on semantic violations than controls (*β* = −2.49, *SE* = 0.376, *χ2* = 30.512, *p* < .001; see Appendix Fig. A1).

### Recognition task

#### Behavioral performance

Mean accuracy in the recognition task was at 63% (*SD* = 9, see Fig. 3A), thus leaving enough variance that could potentially by explained by conditions and/or P600 amplitude in the preceding encoding task (sentence comprehension, see next section). Participants were significantly better in correctly identifying unseen words (*M* = 80%, *SD* = 12) than seen words (*M* = 57%, *SD* = 12; *β* = −1.385, *SE* = 0.205, *z* = −6.741, *χ2* = 30.429, *p* < .001)^4^. Within seen words, words that were previously presented in syntactic violation sentences were remembered significantly worse (*M* = 50%, *SD* = 12) than control sentence words (*M* = 58%, *SD* = 14; *β* = −0.282, *SE* = 0.059, *z* = −4.781, *χ2* = 22.837, *p* < .001). In contrast, words that were previously presented in semantic violation sentences were remembered significantly better (*M* = 63%, *SD* = 14) than control sentence words (*β* = 0.307, *SE* = 0.079, *z* = 3.877, *χ2* = 12.845, *p* < .001).

**Figure 3.**
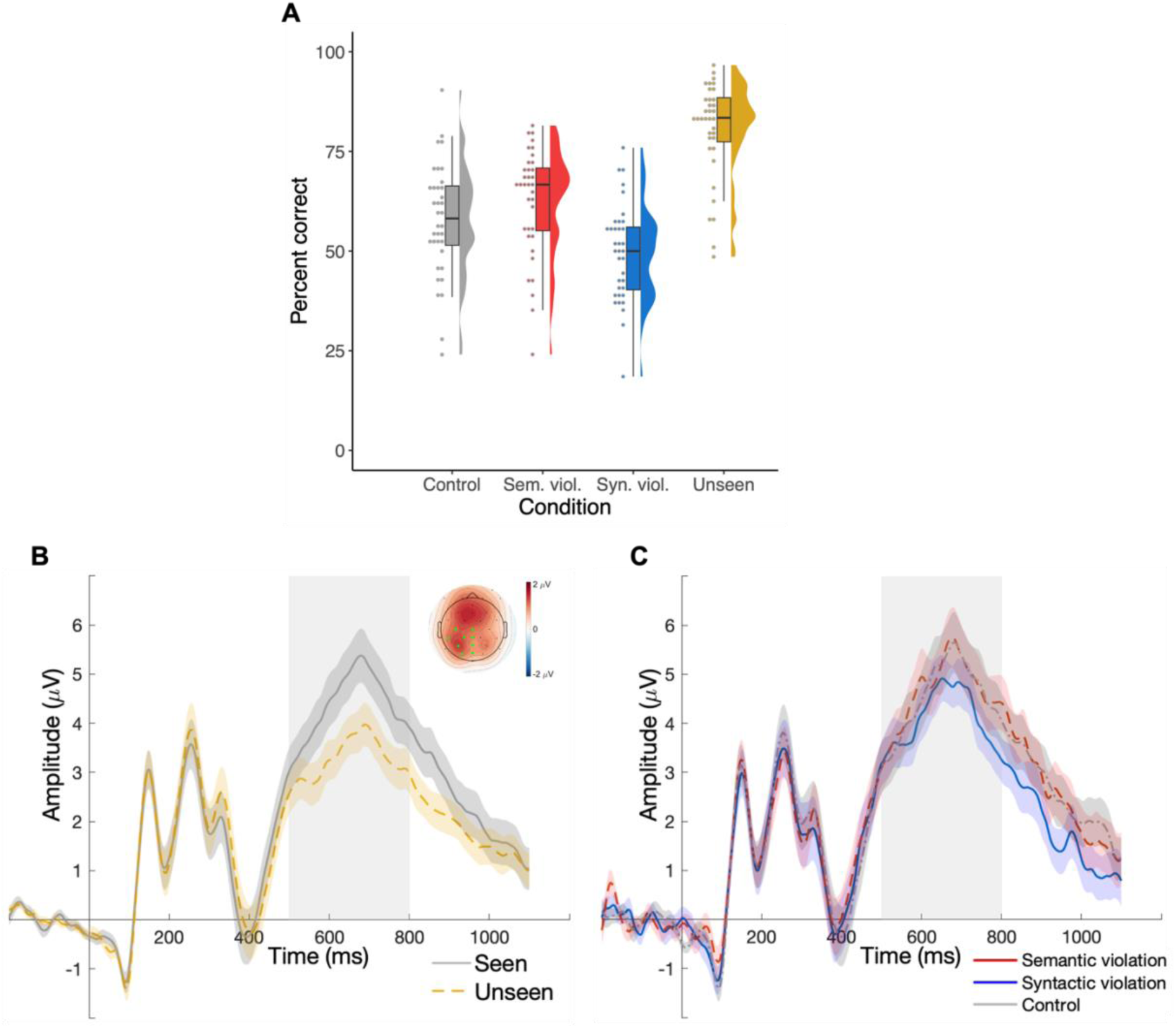
Recognition task results. (**A)** Recognition performance in each of the four conditions. (**B)** Old/new recollection ERP effect: Grand averaged ERP amplitude for the two conditions of the recognition task, time-locked to the onset of the word within the posterior ROI channel cluster (marked in light green on the topographic map). Plot is based on the subset of the data with correct responses (hits and correctly identified new words). Trial-wise ERP amplitudes were calculated as the mean in the time interval marked by the grey box (500-800ms). The topographic map shows the mean amplitude difference between seen and unseen words within the same time window. Note that “seen” words (grey) encompass seen words of all encoding conditions (controls, semantic violations, syntactic violations). (**C)** Grand averaged ERP amplitudes for the three conditions within the “seen” (i.e., words which had been presented within a control, semantic violation, or syntactic violation sentence in the preceding sentence processing task), time-locked to the onset of the word within the posterior ROI channel cluster. The conditions “semantic” and “syntactic” violations as well as “controls” within seen words in plot A, and C refer to the conditions in which the respective word appeared during encoding (sentence processing).

Additional, exploratory analyses showed that the “discrimination index” quantifying corrected recognition (Pr = p(hit) – p(false alarms), Snodgrass & Corwin, 1988) was moderate (*M* = .39, *SD* = .15)^5^. All but one participant (-.03) had positive scores of at least .13. A one-way ANOVA revealed a significant difference of discrimination index between at least two seen word types (F(2, 105) = [6.337], *p* = .003). Tukey’s HSD Test for multiple comparisons revealed the same pattern as above, namely, that the discrimination index was significantly worse for words that were previously presented in syntactic violation sentences (*M* = .3, *SD* = .14) than words that were previously presented in semantic violation sentences (*M* = .46, *SD* = .18, *p* = .002, 95% C.I. = [−22.3, −4.43]). There was no statistically significant difference between control words (*M* = .39, *SD* = .16) and syntactic violation words (*p* = .221) or between control words and semantic violation words (*p* = .148). Relatedly, the response bias (Br = p(false alarms)/(1-Pr), Snodgrass & Corwin, 1988) was above the threshold .5 in all conditions (overall: M = .3, SD = .15; semantic violations: M = .34, SD = .16; syntactic violations: M = .27, SD = .14; controls: M = .3, SD = .16) indicating a conservative response bias. A one-way ANOVA revealed no significant differences of response bias between seen word types (F(2, 105) = [1.966], *p* = .145).

#### Parietal old/new recollection ERP effect

Next, we investigated whether there are neural markers of recognition. We found a classic parietal old/new recollection effect in that ERP amplitudes between 500-800 ms were significantly larger in response to correctly identified seen compared to unseen words (*β* = 1.073, *SE* = 0.254, *χ2* = 15.335, *p* < .001; see Fig. 3B). Note that although the effect was significant within our left-parietal ROI, the topographic plot in Figure 3B indicates that the effect was strongest in mid-frontal regions. Within seen words (see Fig. 3C), there was no amplitude difference during recognition between words previously presented in a control sentence and words previously presented in a syntactic violation sentence (*β* = −0.303, *SE* = 0.289, *χ2* = 1.099, *p* = .295) or semantic violation sentence (*β* = 0.086, *SE* = 0.298, *χ2* = 0.087, *p* = .769).

#### Familiarity ERP effect

As stated in our preregistration, we explored whether the familiarity ERP (frontal negativity, FN400) would also be affected by our manipulation. However, there was no effect of word type on the FN400: Mid-frontal amplitudes between 300-500 ms did not significantly differ between seen and unseen words (*β* = 0.336, *SE* = 0.275, *χ2* = 1.498, *p* = .221; see Appendix Fig. A2). We thus refrained from further analyses relating the FN400 during recognition to the P600 during encoding.

### Relation between P600 during encoding and measures of recognition

Here, we combined the data from the two tasks to investigate whether the P600 during sentence processing (encoding) predicts measures of recognition in the old/new decision task (see Fig. 4). “P600” was defined as the single-trial ERP response to violations during encoding, thus analyses in this section are based on trials in the violation conditions only. Note that for exploratory purposes, we conducted the same analyses regarding the N400 during encoding, but the negativity was not predictive of any of the measures of recognition (see Appendix for detailed results).

**Figure 4.**
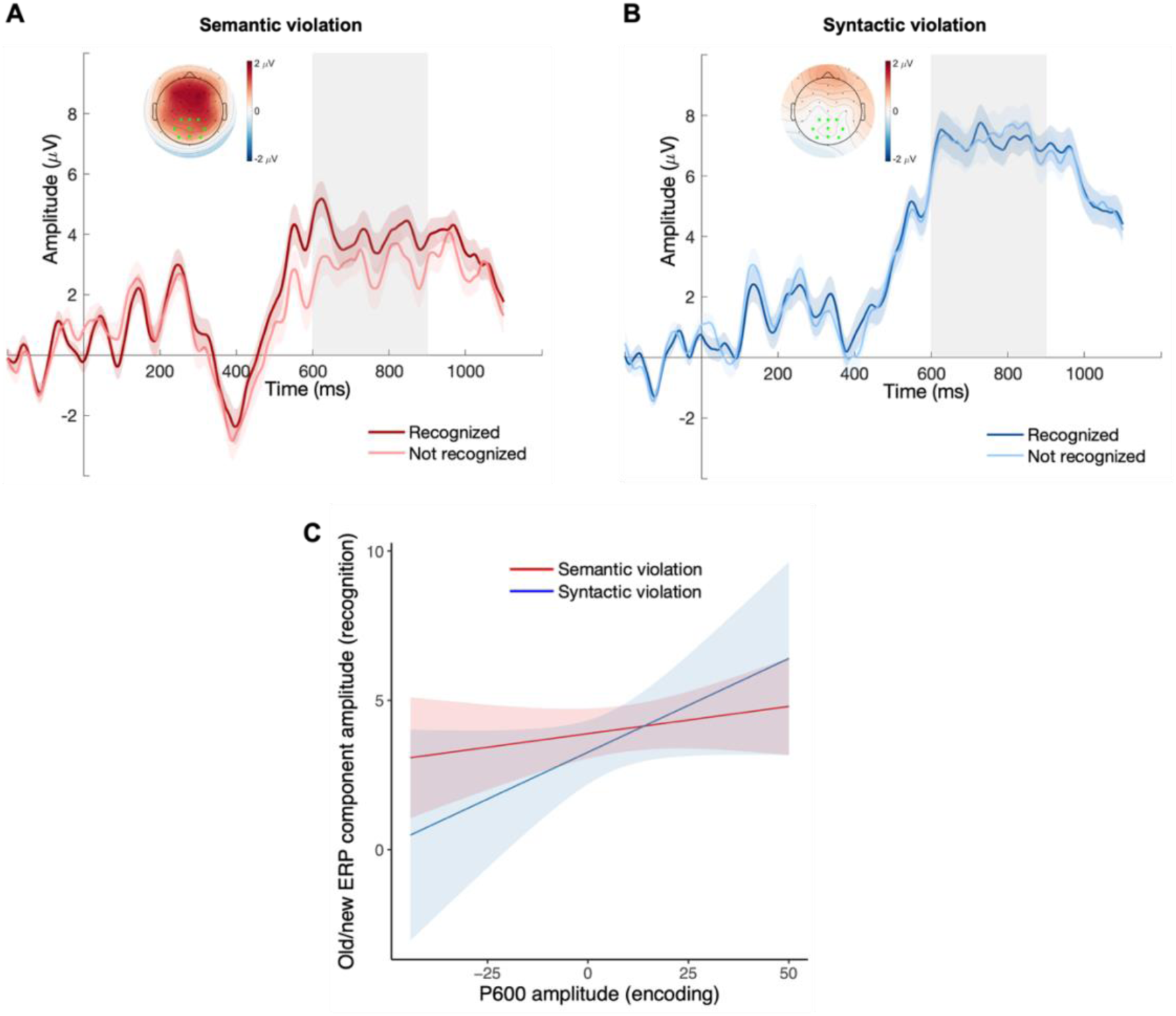
Relation between P600 during sentence processing (encoding) and measures of explicit memory of the respective target noun in the recognition task. Top panel: P600 during encoding (sentence processing) split by recognition performance, separate for (**A**) semantic and (**B**) syntactic violations. Error bands indicate the standard error of the mean (SEM). The topographic maps show the mean amplitude difference between words recognized and not recognized within the P600 time window. Bottom (**C**): Mixed effects model output predictions for the relationship between P600 amplitude at encoding and recognition ERP amplitude (i.e., within temporal and spatial ROI of old/new recollection effect) during recognition, split by violation type. Shaded areas in C display 95% confidence intervals. In all plots, the two violation types “semantic” and “syntactic” refer to the conditions in which the respective word appeared during encoding (sentence processing).

#### P600 and recognition performance

We first investigated whether the parietal P600 amplitude in the encoding task generally predicted performance in the recognition task. This logistic model did not reveal any relationship between the P600 and recognition performance overall (*β* = 0.0003, *SE* = .004, *z* = 0.067, *χ2* = 0.004, *p* = .947). However, there was a significant interaction effect between P600 and condition on recognition performance (*β* = 0.019, *SE* = 0.009, *z* = 2.078, *χ2* = 4.314, *p* = .038). This interaction showed a positive relationship between the P600 and recognition performance in the semantic violation condition (*β* = 0.016, *SE* = 0.007, *p* = .014, Fig. 4A) but not in the syntactic violation condition (*β* = −0.003, *SE* = 0.006, *p* = .699, Fig. 4B). Thus, if sentences during encoding contained a semantic violation, the amplitude of the respective parietal P600 positively predicted whether the target word would be recognized later on. Subsequent, exploratory models also confirmed that this effect was also present in the left-frontal cluster (*β* = 0.025, *SE* = 0.008, *z* = 3.281, *χ2* = 10.918, *p* < .001), where we found the semantic P600 in the first place (see above). Lastly, we explored whether a violation was necessary for the memory effect, and indeed P600 amplitudes in the control condition did not significantly predict recognition performance (*β* = 0.004, *SE* = 0.005, *z* = 0.744, *χ2* = 0.549, *p* = .459). As one would expect, P600 amplitude variance was low in the first place, significantly lower than in sentences containing violations (*F*(1,6613) = 41.505, *p* < .001).

#### P600 and parietal old/new recollection component

Additionally, we were interested in whether the P600 during encoding was predictive of a neural marker of later recognition, namely the amplitude of the parietal old/new recollection ERP component. To that end, we regressed the trial-wise amplitude of the P600 component in response to sentence target words during encoding against the trial-wise amplitude of the old/new recollection component during recognition of the respective old words^6^. Indeed, the model revealed a positive relationship between the two components: The larger the P600 in the posterior cluster during encoding, the larger the amplitude of the old/new recollection component during recognition (*β* = 0.038, *SE* = 0.018, *χ2* = 4.58, *p* = .032; see Fig. 4C). The additional model including condition as an interaction factor revealed that this relationship was not dependent on the type of violation (*β* = −0.045, *SE* = 0.029, *χ2* = 2.316, *p* = .128). We only found a positive trend between the amplitude of the old/new recollection component (measured at the usual left-parietal cluster) and the semantic P600 in the left-frontal cluster (*β* = 0.049, *SE* = 0.03, *χ2* = 2.718, *p* = .099).

## Discussion

The present study investigated a potential link between the P600 during sentence reading and measures of later explicit memory using an old/new word recognition task. During encoding, we found a parietal P600 in response to both syntactic violations and semantic deviants (henceforth: syntactic P600 and semantic P600, respectively), while the latter induced additional frontal activations. In the subsequent word recognition task, seen targets that had appeared as a semantic violation during encoding were recognized better, but those that had appeared within syntactic violations were recognized worse than correct control targets. In addition, we found an old/new recollection ERP effect in that correctly identified seen targets generally elicited more positive amplitudes as compared to previously unseen targets at parietal sites. Crucially, the P600 amplitude during encoding (sentence comprehension) was related to these recognition effects in the subsequent memory task on a trial-by-trial basis: Recognition performance was better for violated words that had previously elicited a larger P600 as a semantically deviant word during sentence comprehension, but not for (previous) syntactic violations. Moreover, the amplitude of the old/new recollection ERP component during recognition of seen targets was positively related to the amplitude of the P600 during encoding. This relationship of ERPs between encoding and recognition was present for both semantic deviants and syntactic violations.

In sum, we find that the P600 predicts later recognition memory both on the behavioral and neural level. Our results thus demonstrate that the memory effect previously observed regarding the P3 at different latencies can be extended to the P600. Specifically, we replicate the effect that the positivity in response to semantic deviance during encoding predicts recognition performance (Neville et al., 1986), but we do so in a canonical sentence paradigm, thus explicitly linking the memory effects also to the “classic” P600. In addition, our study contributes the novel finding that the P600 during encoding predicts neural markers of recognition (the amplitude of the old/new recollection component), an effect that – to our knowledge – no previous study has tested with respect to any late positivity during encoding. This relationship to this neural marker was present not only for the semantic P600 but also for the syntactic P600, thereby for the first time also linking structural (syntactic) violations to later explicit memory. Note that although also a “late positivity”, the amplitude of the old/new recollection ERP component is sensitive to the amount of information recollected from a previous event (see e.g., Curran et al., 2006; Rugg & Curran, 2007) and not to any linguistic violation or saliency. Thus, the relationship between the P600 during encoding and the old/new recollection ERP component at recognition cannot simply be explained as a positive relationship between the initial P600 and a second P600 upon repeated presentation of the target word. Such explicit memory effects further link the late, positive P600 component during language comprehension to the P3 and the LC/NE system, suggesting a more domain-general nature of the component. The link between the P600 and measures of explicit memory indicates that the neurocognitive processes that deal with salient and anomalous aspects in the linguistic input in the moment will also be involved in keeping this event available for later recognition.

We replicated the predictive relationship between the late positivity during encoding and subsequent memory in semantic contexts (e.g., Friedman & Trott, 2000; Höltje et al., 2019; Hubbard & Federmeier, 2024; Neville et al., 1986; Paller et al., 1987, 1988). Interestingly, we found this subsequent memory effect specifically within semantic violations, that is, when the target word was semantically incongruent (implausible) in the sentence, whereas in several of the aforementioned studies (e.g., most recently Höltje et al., 2019), this effect was present in the semantically congruent condition or both. Notably, what also differs is that in our study, semantically incongruent words were remembered better than congruent words, whereas the opposite has often been the case previously (e.g., Höltje et al., 2019; Neville et al., 1986; Van Kersteren et al., 2013; DeWitt et al., 2012; but see Corley et al., 2007; Federmeier et al., 2007; McFalls & Schwanenflugel, 2002). Thus, several previous studies found a memory advantage for semantically congruent (vs. incongruent) words and a subsequent ERP memory effect specifically for these congruent words, a pattern that is entirely reversed in the present study. What might come into play here is distinctiveness, as memory for semantically incongruent items is better when their proportion is low than when their proportion is higher (Reggev et al., 2018). Indeed, whereas a common congruency manipulation in memory studies has equal numbers of items in each condition, semantic violations only made up 22% of our sentences, as typical for P600 studies as the ERP effect reduces with more frequent anomalies (Yano et al., 2021). In addition, previous memory literature has often used simple categorization frames (e.g., “A four-footed animal. Dog / Sapphire”, Höltje et al., 2019), which might not initialize a situational model and semantic expectations quite like sentences. Our findings align well with a very recent set of studies by Hubbard and Federmeier (2024) who also used sentences with a relatively lower proportion of incongruent (implausible) endings (33%). Just as in our study here, memory for incongruent endings was superior to congruent endings and P600 amplitudes elicited by these incongruent endings during sentence reading related to successful recognition of those words. Interestingly, they also found that sentences themselves were less likely recalled when they included an incongruent compared to a congruent ending, indicating that encoding of the context suffers when an implausible word does not fit in. Thus, in 50:50 categorization tasks, schema incongruent exemplars might be discarded quickly, leading to a weak memory trace. A lower proportion of semantically incongruent target words in sentences however might draw attention to these words specifically, which elicits the P600 and benefits memory, though trading in the memory for the sentence context itself.

In our study, the role of the P600 in memory depended strongly on the type of violation. In particular, only the amplitude of the semantic, but not syntactic P600 predicted later behavioral recognition (performance), but both explained the neural marker of recognition (i.e., amplitude of the old/new ERP component). There are at least two possible explanations for this dissociation. Perhaps this is a statistical issue: A ceiling effect of the syntactic, but not semantic P600 could diminish much of the variance within the amplitude that could reliably serve as a predictor for a binary outcome (correctly vs. incorrectly recognized). However, our data revealed that variances of the syntactic and semantic P600 amplitudes did not significantly differ from each other (*F*(1,70) = 0.009, *p* = .924), making this explanation unlikely to account for the different results patterns regarding the two violation types. Alternatively, this dissociation could be due to actual cognitive processing differences. Our data suggest that the relationship between the P600 amplitude during encoding and explicit memory was stronger for the semantic deviances than syntactic violations. This could be due to differences in the formed memory representation, depending on the respective salient aspect: Semantic mismatches might form a memory trace for the word form and meaning of the nouns themselves (which was exactly what was tested in our recognition task, see below) whereas syntactic violations possibly elicit a memory trace specific to the article-noun mismatch and possibly only a weak trace of the noun itself. A “deeper” semantic processing (and thus a stronger semantic memory representation) during encoding might have led to an actual behavioral recognition advantage in our target word recognition task. This possibility is also supported by our finding that recognition performance was generally better for semantic violations than controls. In comparison, performance was worse for target words previously involved in syntactic violations (than controls), suggesting that these types of violations even distracted from the meaning/form of the involved nouns. In contrast to this discrepancy in behavior, the neural marker old/new ERP amplitude was not sensitive to the type of violation during encoding. Assuming that brain responses are more sensitive to subtle processing differences that do not necessarily translate into overt behavior (Luck et al., 1996; Wilkinson & Halligan, 2004), the weak memory trace of the word form elicited by syntactic violations could have been enough to be picked up by the old/new ERP component but not strong enough for a behavioral benefit. Relatedly, the current design created a mismatch between the processes involved in encoding and test for syntactic violations (morphosyntactic mismatch between words vs. single word form and meaning) but not semantic violations. This could also contribute to our pattern of results as parietal subsequent memory effects have been shown to depend on the congruency between encoding and retrieval processes (Bauch & Otten, 2007). Lastly, the salient morphosyntactic mismatch between noun and adjective might be interpreted as a case of an inter-item association, which might not relate to (parietal) subsequent memory effects the way item information in the case of salient semantic violations does (Forester & Kamp, 2023; see also Mecklinger & Kamp, 2023). Further research is required to test these posthoc interpretations, and we are currently following up on these questions by behaviorally probing the memory of each salient aspect specifically.

The observed link between the P600 and explicit memory is in line with the hypothesis that the positivity reflects norepinephrine release from the LC to task-relevant cortices. By activating attentional networks and concurrently projecting to memory-relevant limbic structures, processing the stimulus forms a more robust memory representation (Aston-Jones & Cohen, 2005; Sara, 2015). While the LC/NE framework might not be necessary to explain the memory effects, it is – to our knowledge – the only P600 theory that explicitly predicts a (neurobiologically grounded) link to memory formation. Further, alternative theories that do link late positivities to memory do not necessarily conflict with the role of the LC/NE system. For example, if the P600 is interpreted as a variant of the P3, another theory that would predict such a memory effects is the context updating hypothesis (Donchin, 1981), which proposes that the P3 signals changes in the mental model of the environment in response to unexpected events. However, regarding memory effects, the two theories are not incompatible: While the context updating hypothesis is a theory about the cognitive process, the LC/NE hypothesis pertains to the underlying neurobiological basis. Indeed, Donchin (1981) even explicitly suggests norepinephrine as the neurophysiological system responsible for the memory effects. In the same vein, some cognitive theories of the P600 itself may not necessarily be incompatible with the involvement of the LC/NE system and downstream memory effects (e.g., Brouwer et al., 2012, 2017; Fitz & Chang, 2019; Kuperberg et al., 2020; Li & Ettinger, 2023; van de Meerendonk et al., 2009, 2010) even though they would need to make additional assumptions. It could also be argued that the memory effects are induced by the saliency of the linguistic violations, which thus probably attract attention. Both saliency and attention are generally known to foster memory encoding (e.g., Long et al., 2018; Santangelo, 2015). However, a mechanistic understanding is needed how these factors are connected. Interestingly, neuromodulation via NE is one of the most established theories connecting saliency to attention (neural gain) and memory (Aston-Jones & Cohen, 2005; Bouret & Sara, 2005). Further, the left inferior frontal gyrus (LIFG) has alternatively been proposed as a neural generator of the P600 (Brouwer & Hoeks, 2013). As a major hub for linguistic as well as domain-general processing (e.g., Fedorenko & Thompson-Schill, 2014; Novick et al., 2005; Rogalsky, & Hickok, 2011; Santi and Grodzinsky, 2010) the LIFG would not necessarily be incompatible with concurrent memory effects. Future studies could test the role of the LC/NE system in the connection between the P600 and explicit memory more directly by concurrently measuring NE markers such as pupil dilation (Contier et al., 2024) or manipulating NE via pharmacology or stimulation (e.g., Giraudier et al., 2024; Joseph & Sitaram, 1989; Ventura-Bort et al., 2018).

The current findings also corroborate a proposed dichotomy between the properties and functions of the two ERP components reliably observed during language comprehension: The N400 and P600. It has been suggested that the N400 reflects a more automatic brain response, for instance a variant of a semantic prediction error (e.g., Bornkessel-Schlesewsky & Schlesewsky, 2019; Federmeier, 2007; Kuperberg, 2016; Kuperberg et al., 2020; Rabovsky et al., 2018) or retrieval of words meaning from semantic long-term memory (Brouwer et al., 2012; 2017). In terms of memory, the N400 could then relate to *implicit* learning driving adaptation during language comprehension (Hodapp & Rabovsky, 2021). In contrast, the P600 seems to reflect more controlled aspects of sentence comprehension that require awareness (Batterink & Neville, 2013; Xu, Abdel Rahman, & Sommer, 2019) and attention (Contier et al., 2022; Hahne & Friederici, 1999), which in turn, are associated with more episodic learning (Wagner, 2002). Here, we complementarily demonstrate that these controlled, possibly NE mediated processes reflected in the P600 indeed lead to such *explicit* memory formation. Congruously, the N400 correlates with a neural marker of familiarity in subsequent memory tasks (amplitude of the early old/new ERP component, Meyer et al., 2007), but not with the amplitude of the late, recollection-based old/new recollection ERP component which – conversely – relates to the P600 as we show here. Future research could further elucidate how these different learning signals interact during language comprehension both on the behavioral and neural level, possibly using more naturalistic stimuli.

In conclusion, we find that both the syntactic and semantic P600 during sentence reading positively predict markers of recognition memory. These links to explicit memory mirror those observed for the domain general P3 and are in line with the notion that the P600 is connected to the neuromodulatory LC/NE system in response to salient events. While memory effects of the semantic P600 were straight forward both behaviorally and neurally, our behavioral task might have not allowed us to sufficiently test memory traces induced by syntactic violations, possibly explaining why the respective results diverged on the behavioral vs. neural level. The pattern of results inspires further research on the nature of different memory representations linked to the P600, the question whether these links to memory are causally connected to the neuromodulatory system as well as how they interact with other learning signals during language comprehension.

## Acknowledgements

We thank Lok Yan Lam, Julia Kirschner, and Lioba Berndt for their help with stimulus preparation and/or data collection. Funding was provided by the Research Focus Cognitive Sciences of the University of Potsdam and the German Research Foundation (grant RA 2715/2-1) to Milena Rabovsky.

## Appendix

### Recognition response time analyses

As preregistered, we also analysed recognition response times (RTs) as a potential measure of recognition confidence (Kiani et al., 2014; Volkmann, 1934). Note here again, though, that the recognition task was not speeded, making RTs as a measure of certainty or processing rather unreliable.

#### Effects of condition on recognition response times

Response times (RTs) of correct responses did not differ between seen and unseen words (*β* = −0.028, *SE* = 0.022, *χ2* = 1.538, *p* = .215). Within seen words, RTs were slower on words previously presented in a syntactic violation sentence than words previously presented in a control sentence (*β* = 0.056, *SE* = 0.016, *χ2* = 11.821, *p* < .001). There was no such difference between words from control and semantic violation sentences (*β* = −0.013, *SE* = 0.016, *χ2* = 0.668, *p* = .414).

#### Relation between P600 during encoding and recognition response times

We also investigated whether the parietal P600 amplitude in the encoding task generally predicted confidence in correct responses in the recognition task, quantified by RT. However, we did not find any evidence on a relationship between the P600 and recognition RT overall (*β* = −8.219e-05, *SE* = 1.002e-03, *χ2* = 0.005, *p* = .943). There was no significant interaction between P600 and condition, indicating that this null effect of P600 amplitude was present in both conditions (*β* = 0.0001, *SE* = 0.002, *χ2* = 0.004, *p* = .948; see Fig. 4C). In the same vein, there was no effect of P600 amplitude during semantic violations in the left-frontal cluster on recognition RT (*β* = −0.001, *SE* = 0.001, *χ2* = 0.192, *p* = .661). In conclusion, if sentences during encoding contained a violation, the amplitude of the respective parietal P600 did not predict how fast (i.e., confidently) the target word would be recognized later on.

### Relation between N400 and measures of recognition

To test the specificity of the link to memory for the P600, we additionally investigated whether the same link to memory can be found regarding the N400 in response to semantic violations during encoding (see Fig. A1). Note that these analyses regarding the N400 and its relation to measures of explicit memory were not preregistered.

**Figure A1.**
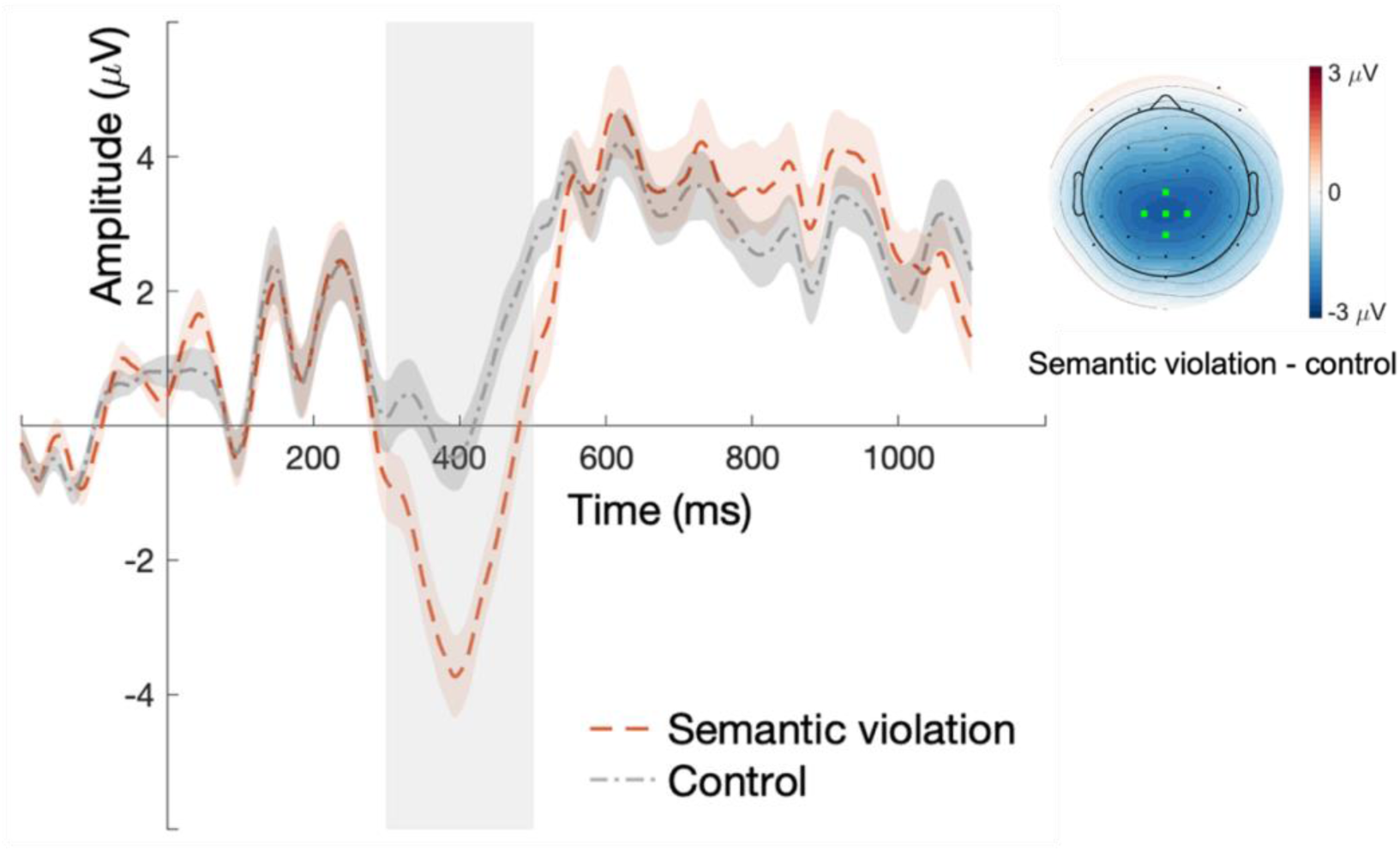
Grand averaged ERP amplitude for the control and semantic violation condition in the sentence processing task at the centro-parietal channel cluster used to measure the N400 (marked in light green on the topographic map). Trial-wise N400 amplitudes were calculated as the mean in the time interval marked by the grey box (300-500 ms). The topographic map shows the mean amplitude difference between semantic violations and controls within the same time window. Error bands indicate the SEM. For statistical analyses of the N400, see main text.

#### N400 and recognition performance

The mixed, logistic model including semantic violation trials in the encoding task and the corresponding recognition trials did not reveal any relationship between the N400 during encoding and recognition performance (i.e., accuracy, *β* = 0.011, *SE* = 0.008, *z* = 1.244, *χ2* = 1.485, *p* = .223). Thus, we did not find any evidence that the amplitude of the N400 on semantic violations sentences during encoding predicts whether the target word would be recognized later on.

#### N400 and recognition confidence

Analogously, we investigated whether the N400 amplitude in the encoding task predicted confidence in correct responses in the recognition task, quantified by RT. However, we did not find any evidence on a relationship between the N400 and recognition RT (*β* = 0.0004, *SE* = 0.001, *t* = 0.354, *χ2* = 0.121, *p* = .728).

#### N400 and parietal old/new recollection ERP effect

Lastly, we also tested whether the N400 during encoding was predictive of the amplitude of the parietal old/new recollection ERP effect during recognition. However, we did not find any evidence of a relationship between the amplitude of the two components (*β* = 0.029, *SE* = 0.024, *t* = 1.193, *χ2* = 1.432, *p* = .231).

### FN400

**Figure A2.**
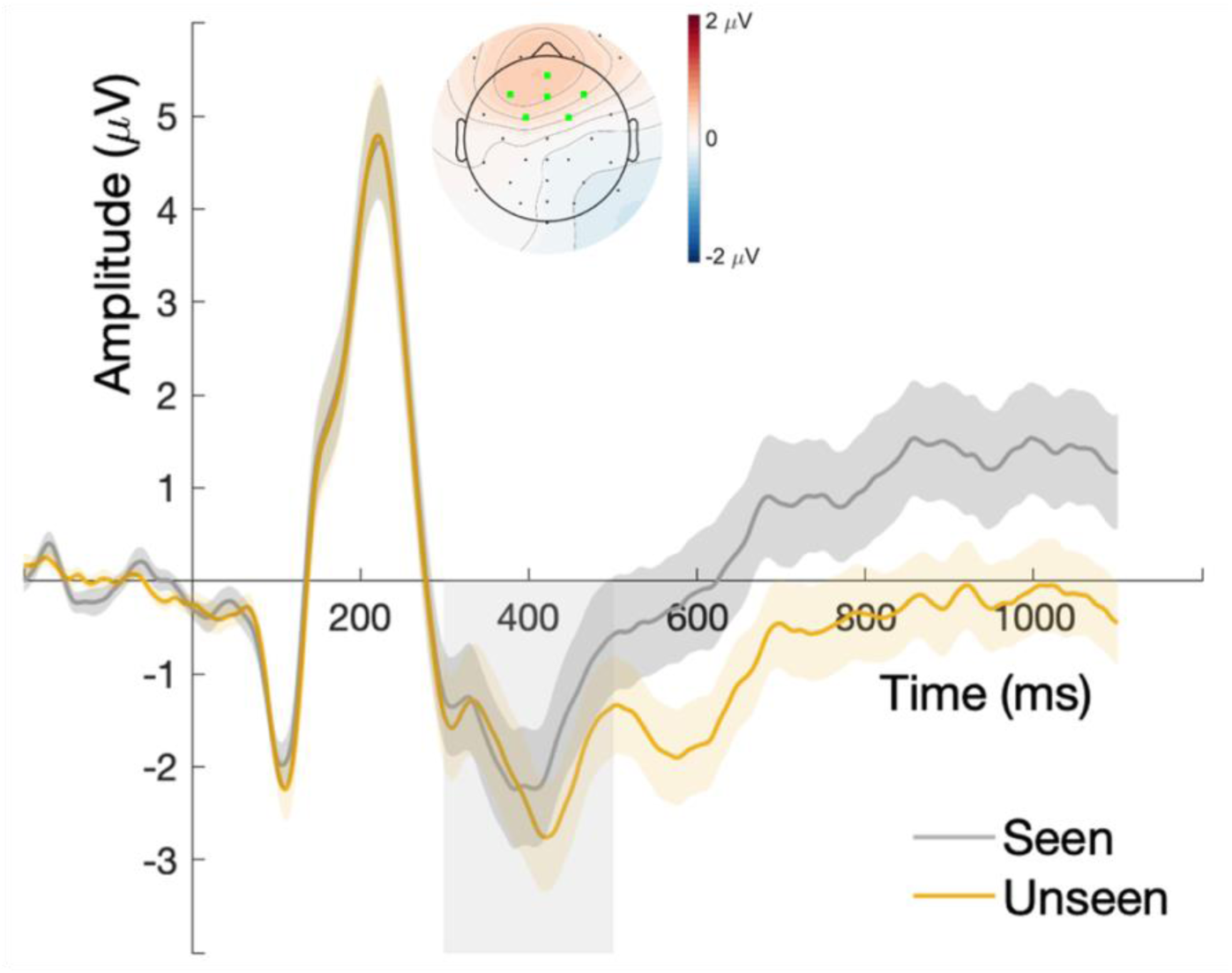
Grand averaged ERP amplitude for the seen and unseen condition in the recognition task at the frontal channel cluster used to measure the familiarity negativity FN400 (marked in light green on the topographic map). Trial-wise FN400 amplitudes were calculated as the mean in the time interval marked by the grey box (300-500 ms). The topographic map shows the mean amplitude difference between seen and unseen words within the same time window. Error bands indicate the SEM. For statistical analyses of the FN400, see main text.

## Footnotes

1 Note that we also planned (i.e., preregistered) to analyze response times as a potential measure of recognition confidence (Kiani et al., 2014; Volkmann, 1934). However, since our focus here was on *explicit* memory and recognition accuracy, decisions on recognition were not speeded and response times might thus not be as informative of memory retrieval as in, for example, implicit memory paradigms (e.g., Hodapp & Rabovsky, 2021). We still report recognition response time analyses in the appendix.

2 Sample size was set to be larger than many previous relevant studies on the P600 (e.g., Friederici et al., 1993; Gouvea et al., 2010; Kuperberg, Sitnikova, et al., 2003; Osterhout & Mobley, 1995; Sassenhagen & Fiebach, 2019; Xu et al., 2021), the P3 and explicit memory effects (e.g., Dunn et al., 1998; Kamp et al., 2013, 2015, 2012; Karis et al., 1984; Sanquist et al., 1980), and late positivities and memory effects in semantic/linguistic contexts (e.g., Friedman & Trott, 2000; Neville et al., 1986; Paller et al., 1987, 1988).

3 We used length, annotated, normalized type frequency, and orthographic neighborhood size (Coltheart) according to *dlexDB* (www.dlexdb.de).

4 Note that we had preregistered this model comparing performance of seen vs. unseen words to include frequency as covariate to account for such systematic differences since it was very unlikely to achieve equal frequency across these two word types. However, as described in the method section, our final words sets for unseen words did not differ significantly in frequency, hence the model was built without frequency as a covariate.

5 Recognition performance both in terms of proportion of correct “old” words and discrimination index was similar to previous studies with comparable stimuli and procedure (e.g., Haeuser & Kray, 2022; Höltje et al., 2019; Mecklinger & Höltje, 2022; Meyer et al., 2007; Rommers & Federmeier, 2018)

6 Please note that even though we adhere to the traditional name “old/new recollection” component, in our analyses we did not focus on the difference between old and new words, but just on single trial amplitudes to old words as new words could not be related to P600 amplitudes during encoding.

